# Mechanism underlying starvation-dependent modulation of olfactory behavior in *Drosophila* larva

**DOI:** 10.1101/781682

**Authors:** Eryn Slankster, Sai Kollala, Dominique Baria, Brianna Dailey-Krempel, Roshni Jain, Seth R. Odell, Dennis Mathew

## Abstract

Starvation enhances olfactory sensitivity that encourage animals to search for food. The molecular mechanisms that enable sensory neurons to remain flexible and adapt to a particular internal state remain poorly understood. Here, we study the roles of GABA and insulin signaling in starvation-dependent modulation of olfactory sensory neuron (OSN) function in the *Drosophila* larva. We show that GABA_B_-receptor and insulin-receptor are necessary for OSN modulation. Using a novel OSN-specific gene expression analysis, we explore downstream targets of insulin signaling in OSNs. Our results strongly suggest that insulin and GABA signaling pathways interact within OSNs and modulate OSN function by impacting olfactory information processing and neurotransmission. We further show that manipulating these signaling pathways specifically in the OSNs impact larval feeding behavior and its body weight. These results challenge the prevailing model of OSN modulation and highlight opportunities to better understand OSN modulation mechanisms and their relationship to animal physiology.

## INTRODUCTION

Starvation increases olfactory sensitivity that enhances an animal’s search for food. This has been shown in insects, worms, and mammals including humans (Cameron et al., 2012; Chao et al., 2004; Ko et al., 2015; Koelega, 1994; Root et al., 2011; Stafford & Welbeck, 2011; J. Tong et al., 2011). However, the mechanisms by which an animal’s starved state modulates sensory neuron function remain poorly understood. Our understanding of these mechanisms significantly improved in the last decade or so from studies that showed how neuromodulators enable changes in the gain of peripheral sensory inputs (Bargmann, 2012; Fields, 2004; Gaudry & Kristan, 2009; Ko et al., 2015). The prevailing mechanistic model for olfactory sensory neuron (OSN) modulation by the animal’s starved state is that during the animal’s starved-state, lower insulin signaling frees production of the short neuropeptide F receptor (sNPFR1), which increases sNPF signaling. Higher sNPF signaling increases presynaptic facilitation of OSNs, which leads to enhanced responses to odors (Bargmann, 2012; Ko et al., 2015; Root et al., 2011). Interestingly, insulin and neuropeptide Y (the mammalian ortholog of *sNPF*) signaling also feature in the vertebrate olfactory bulb (Baskin et al., 1985; Mathieu et al., 2002; Mousley et al., 2006).

However, the above model is incomplete and several questions remain. For instance, the model does not account for the role of GABA signaling, which plays important roles during both starvation and olfactory behavior in flies and mammals (Root et al., 2008; Wan et al., 1997; Weizman et al., 1990). The model also does not account for interactions between GABA and insulin signaling pathways that are known to impact neuromodulation in both fly and mammalian systems: For instance, GABA_B_-Receptor (GABA_B_R) mediates a GABA signal from fly brain interneurons, which may be involved in the inhibitory control of *Drosophila* insulin like peptide (DILP) production (Enell et al. 2010); In mammalian CNS neurons, insulin increases the expression of GABA_A_R on the postsynaptic and dendritic membranes (Sohrabipour et al., 2018; Wan et al., 1997); GABA administration to humans resulted in a significant increase in circulating Insulin levels under both fasting and fed conditions (Li et al., 2015; Sohrabipour et al., 2018). Finally, the model does not account for the ultimate targets of insulin/GABA/sNPF signaling that alter OSN sensitivity to odors and its function.

The above questions are significant because the mechanisms driving neural circuit modulations are fundamental to our understanding of how neural circuits support animal cognition and behavior. If we better understood these mechanisms, we could learn how flexibility and the ability to adapt to a particular internal state are built into the sensory circuit. Understanding the mechanisms by which the starved state of an animal modulates its olfactory sensitivity and thereby controls its food-search behavior is important for both olfactory and appetite research. Finally, we cannot ignore this connection in light of the obesity epidemic and the demonstration that obese adults have reduced olfactory sensitivity (Richardson et al., 2004).

Here, we build upon the prevailing model and argue that GABA and insulin signaling pathways interact within OSNs to mediate starvation-dependent modulation of its function and that defects in these signaling pathways impact larval food-search and feeding behaviors, which in turn impact weight gain. We use the convenient *Drosophila* larval system to demonstrate evidence in support of our argument. Using larval behavior assays, we show that GABA_B_R and insulin receptor (InR) are necessary for starvation dependent increases in larval olfactory behavior. Using a novel OSN-specific gene expression analysis, we show that insulin and GABA signaling pathways interact within OSNs and modulate OSN function by impacting odor reception, olfactory information processing, and neurotransmission. Finally, we show that manipulating these signaling pathways specifically in the OSNs impact larval feeding behavior and its body weight.

## MATERIALS AND METHODS

### Fly strains

Flies were reared on standard cornmeal-dextrose agar food (Nutrifly, Bloomington formulation. Genesee Scientific #66-112) at 25°C and 60% humidity.

*Canton-S* (*CS*) and *w*^*118*^ lines were used as wild type lines in behavioral experiments. The following *OSN-Gal4* drivers were used: *Orco-Gal4* and *O47a-Gal4* (from Dr. John Carlson). The following *UAS-lines* were used: *UAS-InR*^*CA*^ (BDSC #8262), *UAS-InR-RNAi* (VDRC #992), and *UAS-GABA*_*B*_*R2-RNAi* (VDRC #1784). The following *UAS-line* was used for the OSN-nuclei isolation experiments: *UAS-eGFP-MSP300*^*KASH*^ (from Dr. Vikki Weake).

### Behavioral assays

#### Preparation of animals

Behavioral experiments utilized third-instar *Drosophila* larvae (∼96 hours after egg laying). The larvae were extracted from food using a high-density (15%) sucrose (Sigma Aldrich Inc.) solution. Larvae that float to the surface of the sucrose solution were separated into a 1000 ml glass beaker and washed four times with distilled water.

#### Starvation protocol

Washed larvae were allowed to roam freely for 2 hours at RT in a 6 cm petri-dish (Falcon Scientific #351007) containing either 350 μL dH_2_O added to a piece of Kim wipe (starved condition) or 350 μL of 0.2 M sucrose (Acros Organics #177140050) added to a piece of Kim wipe (non-starved condition).

#### Two-Choice behavior assay

The assay was adapted from (Monte et al., 1989) (***Figure 1A***). Larval crawling media were prepared by pouring 10 ml of melted (1.2%) agarose (Genesee Scientific #20-102GP) into 10 cm petri-dishes (Genesee Scientific #32-107). Odor was added to a 6 mm filter disc (GE Healthcare #2017-006) placed on one end of the petri-dish and the diluent, paraffin oil (Sigma-Aldrich, #76235) was added to a filter disc placed on the opposite side. Odor gradients formed remain stable for the duration of the assay (Mathew et al., 2013). Approximately 50 third-instar larvae were placed in the center of the dish and allowed 5 min to disperse in the dark. After 5 min, the number of larvae on each half of the dish were counted to generate the response index (RI), calculated as [eqn. 1].

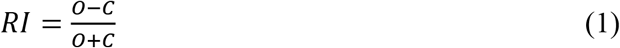

**Figure 1.**
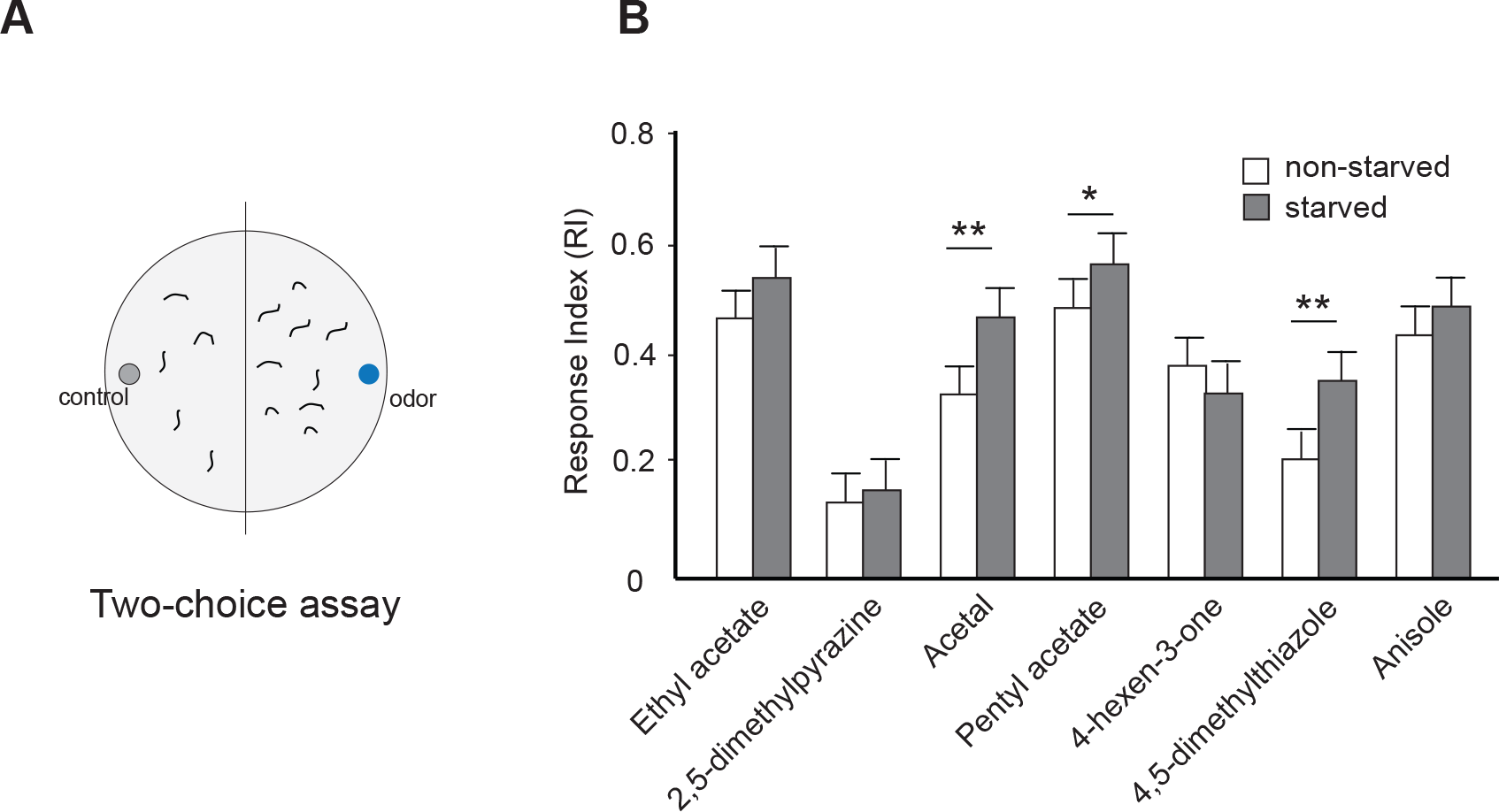
Starvation enhances larval olfactory behavior. (**A**) Schematic for larval two-choice assay. (**B**) Olfactory behavior observed in non-starved (white bars) and starved (grey bars) third-instar *Drosophila* larvae is measured in response to a panel of seven odors on the X-axis. Response index is calculated based on the number of larvae in the odor half and control half and plotted on the Y-axis (n=10 trials). Mean ± SEM. *p<0.05, **p<0.01.

*O* represents the number of larvae that were on the half of the plate containing odorant and *C* is the number on the half containing the control disc. A minimum of ten assays were performed for each odor and condition. Odorants used in these studies were obtained at the highest purity available (≥ 98% purity, Sigma-Aldrich Inc. St. Louis, MO). The temperature of the behavior room was maintained between 22°C and 23°C and between 45% and 50% relative humidity.

#### Feeding behavior assay

The assay was adapted from (Kaun et al., 2007) (***Figure 7A***). Larvae were placed in a 6 cm Petri-dish containing a mixture of 0.2 M sucrose and 0.08% Brilliant Blue R dye (Acros Organics, #191490050). The larvae were allowed to feed on the sugar solution for 15 minutes. After this period, larvae were collected and sacrificed by boiling them for 10 seconds. After rinsing with dH_2_O, larvae were placed on a slide and imaged using a Moticam 10+ microscope camera (Motic). Images were analyzed using Motic Images Plus 3.0 ML. The blue colored area in each larva corresponding to dye intake was measured and normalized against the total area of the larva.

### Immunocytochemistry

Third-instar larval dissection and antibody staining methods were adapted from (Ramachandran & Budnik, 2010) and (Vosshall et al., 2000). Larvae were dissected in Phosphate buffered saline (PBS) and fixed in 4% paraformaldehyde for 30 min at RT. Fixed larval samples were washed three times in PBS and treated with PBS-T (PBS + 0.2% TritonX). Samples were incubated with primary antibodies diluted in 5% BSA for 16-18 hours at 4°C. Following three washes in PBS, samples were incubated with secondary antibodies diluted in 0.4% BSA for 4 hours at RT. Samples were washed again three times, mounted in Vectashield (Vector Laboratories #H-1000) on glass slides and analyzed with a Leica TCS SP8 Confocal Microscope.

InR was stained using a (1:65) dilution of rabbit anti-InR polyclonal antibody (Cloud-Clone Corporation, #PAA895Hu02), which was visualized with a (1:65) dilution of goat anti-rabbit IgG coupled to Alexa Fluor Plus 647 (ThermoFisher, #A32733). To visualize the GABA_B_-Receptor in larval OSNs, we generated a rabbit anti-GABA_B_R1 antibody. To do so, we custom synthesized a 15 amino acid peptide (TVAEAAKMWNLIVLC) specific to the GABA_B_-R1 subunit. This peptide spanning amino acids 121-135 in GABA_B_-R1 was selected because it is a conserved motif across insect species. We used this peptide as an antigen to generate a rabbit polyclonal antibody (Pocono Rabbit Farm & Laboratory Inc.). GABA_B_R was stained using a (1:125) dilution of the rabbit anti-GABA_B_R1 polyclonal antibody, which was visualized with a (1:125) dilution of goat anti-rabbit IgG coupled to Alexa Fluor Plus 647 (ThermoFisher, #A32733). GFP ectopically expressed in OSNs was stained using a (1:125) dilution of chicken anti-GFP polyclonal antibody (Invitrogen, #PA1-9533), which was visualized with a (1:125) dilution of goat anti-chicken IgY coupled to Alexa Fluor 488 (ThermoFisher, #A-11039).

### Gene expression analysis

#### Larval head sample preparation

Starved or non-starved third-instar larvae were used for this preparation. 15 larval heads were dissected for each condition using 3 mm surgical scissors and stored in RNAlater (Invitrogen #AM7020). Samples were homogenized using a mortar and pestle in RLT lysis buffer (Qiagen #74134) before subjecting each sample to RNA extraction protocol.

#### Larval OSN-nuclei sample preparation

This protocol was adapted from (Ma & Weake, 2014) (***Figure 4***). Third-instar larvae of the following genotype: *w*; *Orco-Gal4*; *UAS-GFP-Msp300*^*KASH*^ were used for OSN nuclei isolation. Larvae were placed in PBS during sample collection. A pair of surgical scissors was used to dissect out the dorsal organ of the larvae. Larval mouth hooks provided a visual landmark during the dissection. Incisions were made so as to exclude larval brains from the final sample collection. Dissected samples were stored in cold PBT (PBS plus 0.1% Tween-20 (VWR #0777)). Samples were homogenized using a mortar and pestle. Pre-isolation samples were collected to determine nuclear yield, nuclei integrity, and to determine transcript levels of target genes prior to nuclei isolation. Affinity based isolation of nuclei was performed as described in (Ma & Weake, 2014). Briefly, *GFP-Msp300*^*KASH*^ tagged OSN nuclei were pulled down using a Chicken anti-GFP antibody (Invitrogen #PA1-9533) bound to magnetic Dynabeads™ Protein G (Invitrogen #10003D). Antibody-bound beads and homogenate were placed in a magnetic rack, MagRack6 (GE #28948964) for 2 minutes to allow the magnetic beads to bind to the magnet. Homogenate containing the unbound nuclei fraction was removed (***Figure 4C***). Following 3× washes with a Wash buffer, post-isolation samples were suspended in 350 μl RLT buffer and stored at −20C until they were subjected to the RNA extraction protocol.

#### RNA extraction and cDNA library preparation

For each sample, total RNA was isolated using the RNeasy Plus Mini kit with gEliminator columns (Qiagen #74134). An additional gDNA digestion was conducted using the TURBO DNA-free kit (Invitrogen #AM1907) (larval head samples) or with RNase-Free DNase Set (Qiagen, #79254) (larval OSN nuclei samples). OSN samples were eluted into a total of 60 μL RNase-free water and 1μL RNAsecureTM RNase Inactivation Reagent (Invitrogen #AM7005). RNA Clean & Concentrator-5 (Zymo Research # R1013) was performed according to manufacturer’s protocol and RNA was eluted into 12 μL of RNase-free water. RNA quantification was performed using a Nanodrop 1000 spectrophotometer (Thermo Scientific). Larval head cDNA library was constructed from 1 μg of the total RNA using Superscript VILO Master Mix (Invitrogen # 11755050). Larval OSN nuclei cDNA library was constructed from 100-200 ng RNA with Superscript IV VILO Master Mix (Invitrogen #11754050). All cDNA samples were diluted to the equivalent of 1ng/μl RNA.

#### Relative gene expression analysis

Real time quantitative PCR (RT-qPCR) was performed to compare gene expression differences in head (***Figure 3***) and OSN-nuclei samples (***Figure 5***) of third instar *Drosophila* larvae. The MIQE guidelines were followed as far as possible (Bustin et al., 2009). Primer sequences for individual genes were derived from FlyPrimerBank (flyrnai.org), designed using PrimerBlast, or obtained from literature (***Supplementary Table 1***). Melt curve analyses were performed for each reaction to confirm primer specificity. Standard curves were used to calculate primer efficiency and were performed using a minimum of three serial dilutions of cDNA within an experimentally determined amplifiable range (***Supplementary Table 1***). DNA contamination was checked in all samples using primers that spanned exons of several genes.

A 20 μL RT-qPCR reaction included cDNA template synthesized from 1 ng RNA, 0.4 μM of each primer and SsoAdvanced Universal SYBR Green Supermix (Bio-Rad # 1725270). The RT-qPCR was performed on a CFX96 C1000 Touch™ Real-Time PCR detection system (Bio-Rad) with thermal cycling conditions as follows: an initial denaturation of 95°C for 30 sec, followed by 40 cycles of 95°C for 10 sec and 60°C for 30 sec. Each reaction was conducted in triplicates. Occasional reactions within the triplicates with standard deviation > 0.3 were omitted from analysis as PCR outliers.

To demonstrate enrichment of OSN-specific genes relative to other neural genes in the post-isolation samples (***Figure 4D***), we used expression levels of neural genes: *APPL* and *Nrv2* for normalization. *Nrv2*, *Act42a*, *TBP*, and *EF1* were used as reference genes. This combination was picked because it yielded the lowest stability values among prospective reference genes, including *ELAV*, *Orco*, and *eGFP*. The BIO-RAD CFX software measured the collective reference gene expression stability yielding a mean coefficient variance <0.250 and a mean M value, a measure of reference gene expression stability <0.5. For larval heads samples (***Figure 3***), CV = 0.0966, M=0.2116. For larval OSN samples (***Figure 5***), CV=0.17, M=0.3821. Statistical differences between biological sets were calculated within the software.

### Larval body-weight measurements

To quantify larval body-weight, we collected exactly 100 third-instar *Drosophila* larvae (96 h after egg laying), washed and dried the larvae, and carefully measured their combined weight on a Mettler-Toledo Precision balance. We repeated this process 10 times for each genotype considered.

### Statistical analysis

Statistical analyses were performed using Statistica (Statsoft Inc.) unless otherwise noted. All behavior and gene expression data were expressed as mean ± standard error of the mean. For the two-choice behavioral assays the Shapiro-Wilk test was used to assess the normality of distribution of the calculated RI. The distribution of RI for wild-type third instar larvae for each condition followed normal distributions. For the data in ***Figure 1B***, statistical analysis was performed using a two sample T test comparing the non-starved and starved condition for each odor. A B-Y correction for multiple comparisons was performed. This correction is less conservative than a Bonferroni correction and is defined as [eqn. 2].

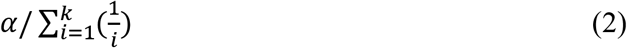

The value for α was 0.05 and k is defined as the number of comparisons (Narum 2006). For the data in ***Figure 2H***, the distribution of the RI for each condition did not follow normal distributions. Analysis was done with the Mann Whitney U test comparing the non-starved and starved conditions for each genotype. A B-Y correction for multiple comparisons was performed. To compare the RI between each genotype for each condition a Kruskal-Wallis multiple comparisons test was performed. The RT-qPCR statistical analysis was performed using the proprietary BIO-RAD CFX software (ver 3.1), which determines mean values and standard deviations and statistical differences were evaluated using t-tests and one-way ANOVA. In all cases, data are presented as relative normalized expression ± SEM (***Figures 3B,C***, ***4D***, ***5B,C***). Expression data were normalized to the expression levels of four genes (*Act42a*, *EF1*, *Nrv2*, and *TBP*). The software helped determine the collective reference gene expression stability by calculating a mean coefficient variance (CV) and a mean M value (M). For both head and OSN-nuclei samples, CV<0.25 and M< 0.5 were obtained. For the larval body weight data (***Figure 6***) and larval food consumption data (***Figure 7B***), boxes represent interquartile ranges. Bars, if shown, represent non-outlier range as defined by Statistica. ANOVA was performed to compare whether the distributions are different from one another.

**Figure 2.**
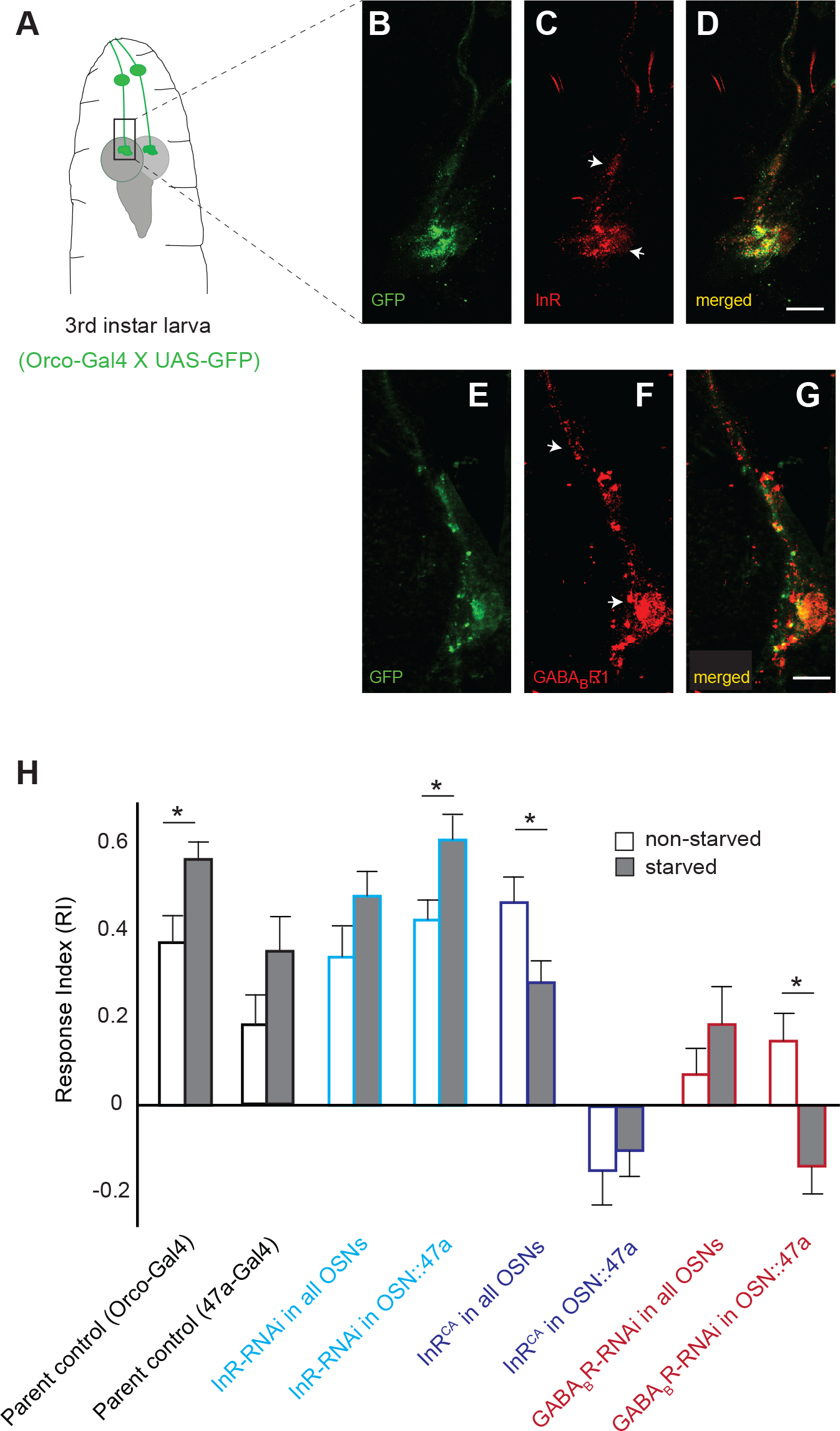
*InR* and *GABA*_*B*_*R* are expressed in larval OSNs. Their expression levels impact starvation-dependent modulation of larval olfactory behavior. (**A**) Cartoon depicting the front end of a third-instar *Drosophila* larva including first-order OSNs (green) projecting into anterior brain regions. Rectangular inset marks the region of interest during confocal imaging. (**B, E**) α-GFP antibody labels the bundle of larval OSNs. (**C, D**) α-InR antibody labels both OSN terminals and axon bundle (white arrowheads). Scale bar = 20 μm. (**F, G**) α-GABA_B_R1 antibody labels both OSN terminals and axon bundle (white arrowheads). Scale bar = 20 μm. (**H**) Olfactory behavior in response to pentyl acetate (10^−2^, vol:vol) is measured using the two-choice assay in control and test genotypes of third-instar *Drosophila* larvae under non-starved (white bars) and starved conditions (grey bars). Mean ± SEM. *p<0.05.

**Figure 3.**
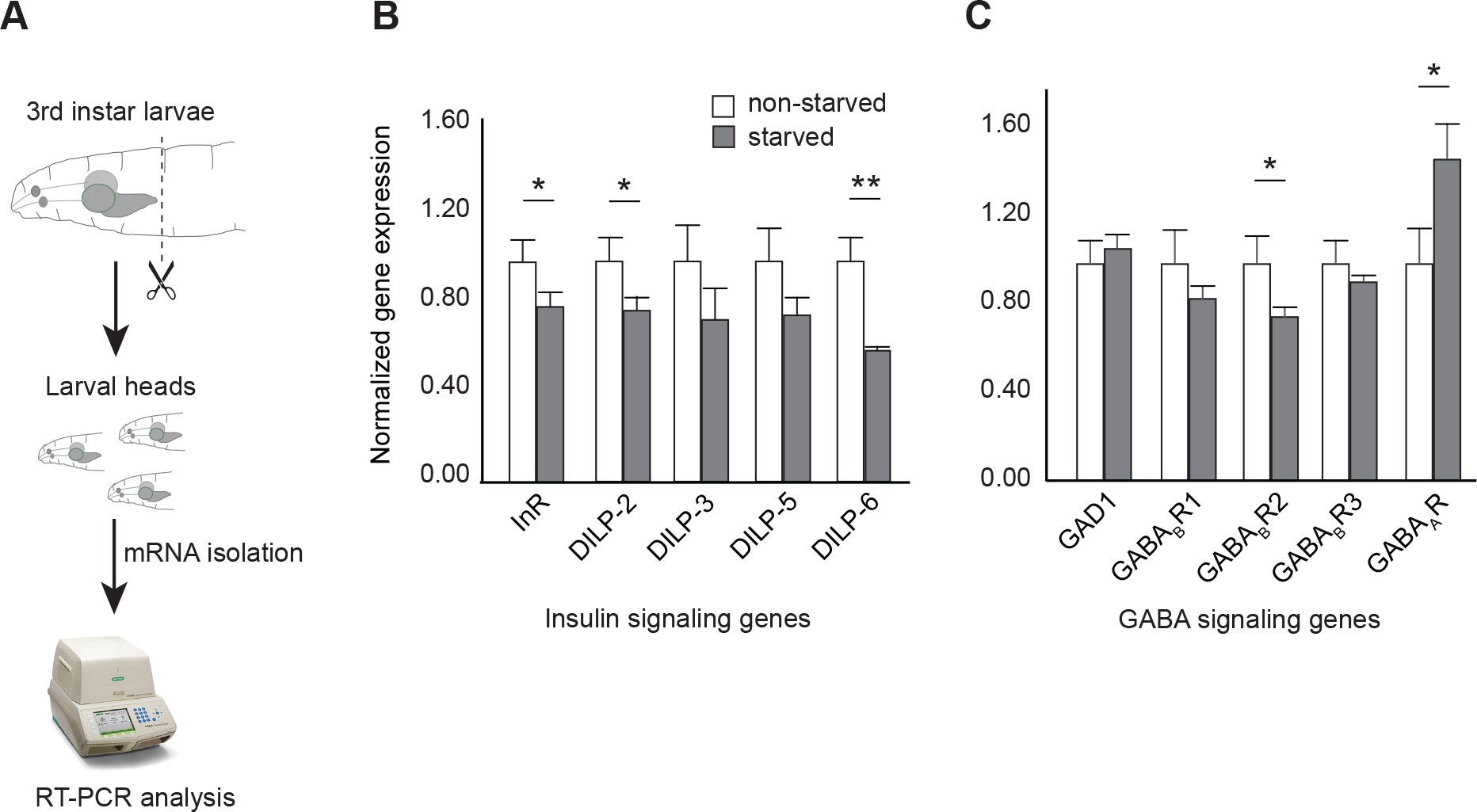
Starvation impacts expression levels of insulin and GABA signaling genes. (**A**) Schematic for gene expression analysis of larval heads. Normalized gene expression of five insulin signaling genes (**B**) and five GABA signaling genes (**C**) are measured following mRNA isolation from non-starved (white bars) or starved (grey bars) larval heads. Mean ± SEM. *p<0.05, **p<0.001.

**Figure 4.**
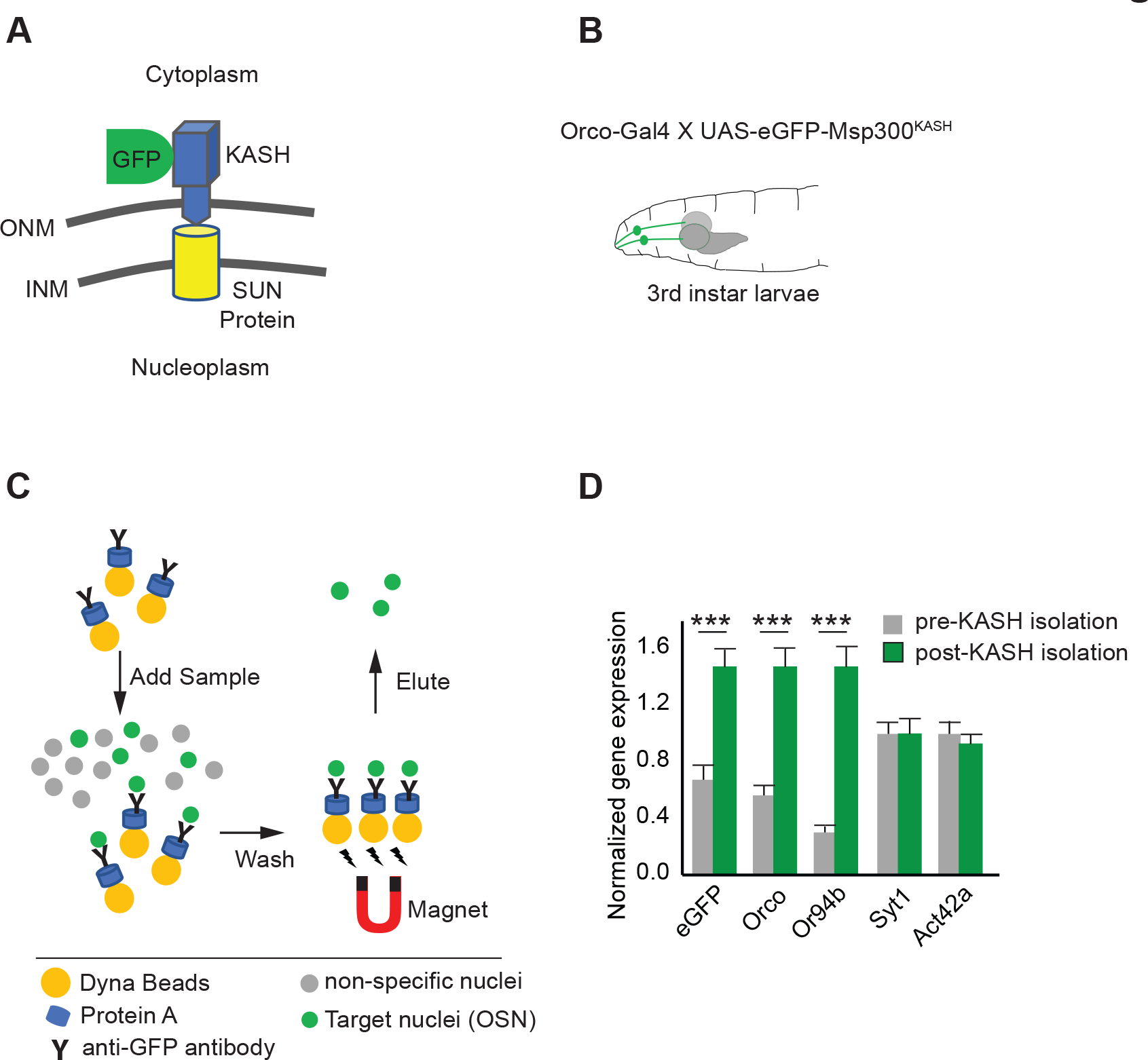
OSN isolation. (**A**) OSN nuclei are genetically tagged with a *eGFP-Msp300*^*KASH*^ that localizes to the nuclear membrane. (**B**) Cartoon showing a third-instar *Drosophila* larva with its two pairs of OSN bundles in green to depict expression of the *eGFP-Msp300*^*KASH*^ construct (**C**) GFP-tagged nuclei (green circles) are separated and purified from other nuclei (grey circles) using a magnetic pull-down method (*see materials and methods*). (**D**) Normalized gene expression levels of OSN specific genes (*Orco, Or94b, eGFP*) and more commonly expressed neural genes (*syt1, Act42a*) are evaluated after RNA purification from pre-isolation and post-isolation OSN-nuclei samples. Mean ± SEM. ***p<0.0001.

**Figure 5.**
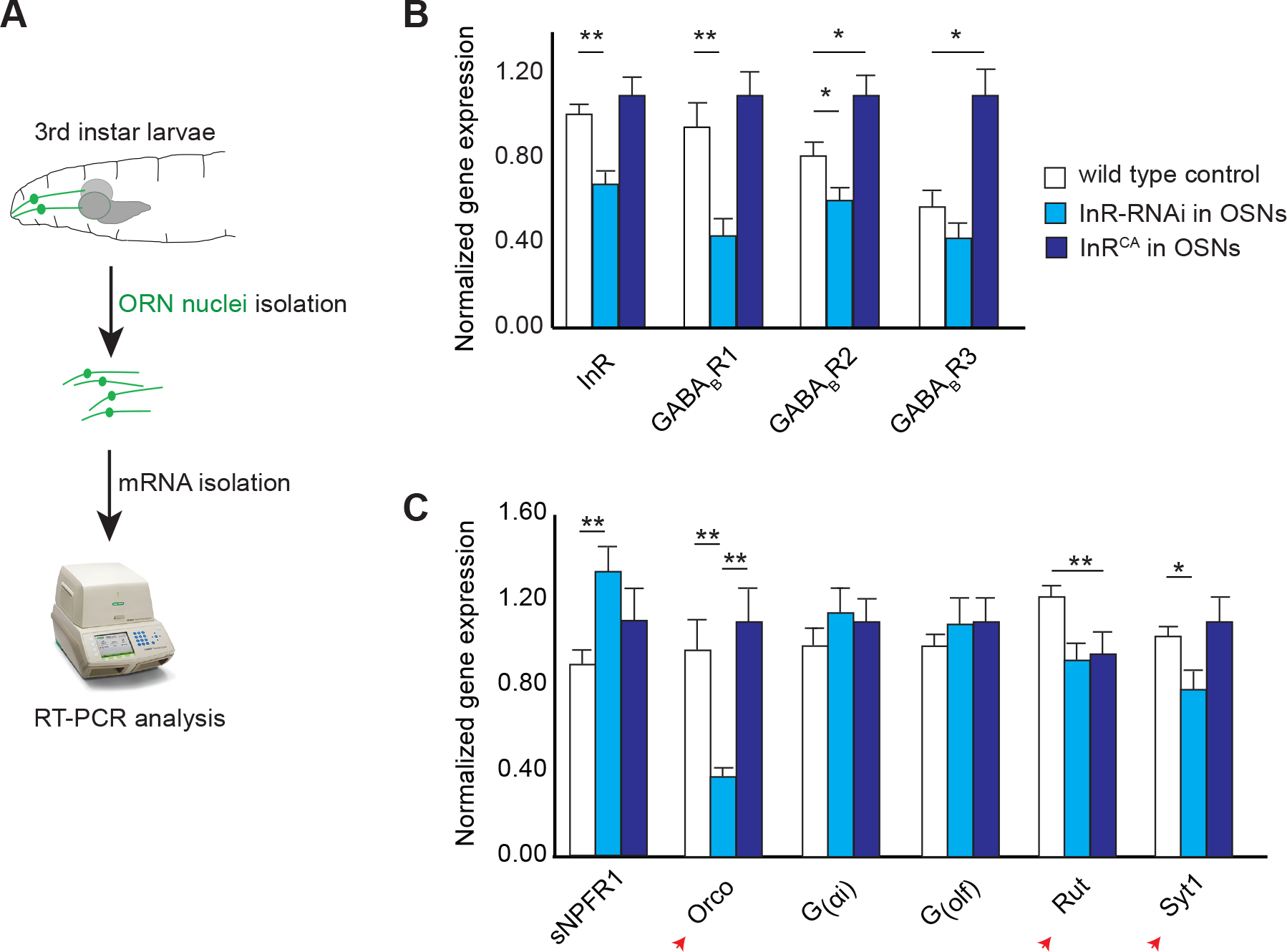
Expression levels of GABA_B_R and other OSN genes are dependent on InR levels. (**A**) Schematic for gene expression analysis of larval OSN nuclei. (**B**) Normalized gene expression levels of *InR* and three *GABA*_*B*_*R*-subunits are measured following OSN isolation and RNA purification from control larvae (white bars) or larvae expressing *InR*-*RNAi* (light blue bars) or *InR*^*CA*^ (dark blue bars) in OSNs. (**C**) Normalized gene expression levels of *sNPFR1*, *Orco*, *G*_*αi*_, *G*_*olf*_, *Rut*, and *Syt1* are measured following OSN isolation and RNA purification from control larvae (white bars) or larvae expressing *InR*-*RNAi* (light blue bars) or *InR*^*CA*^ (dark blue bars) in OSNs. Mean ± SEM. *p<0.05, **p<0.001.

**Figure 6.**
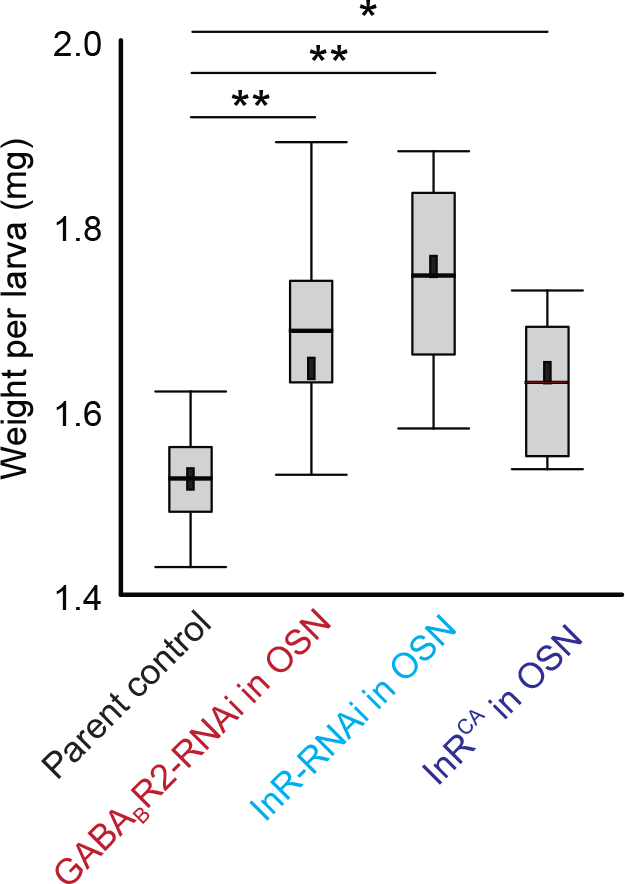
Larval body-weight measurements. Larval body-weight (Y-axis) is measured in control and test genotypes in which GABA_B_R2 or InR levels (X-Axis) are altered in the OSNs. Boxes are interquartile ranges. Bars are the non-outlier range as defined by Statistica software. *p<0.01, **p<0.001.

**Figure 7.**
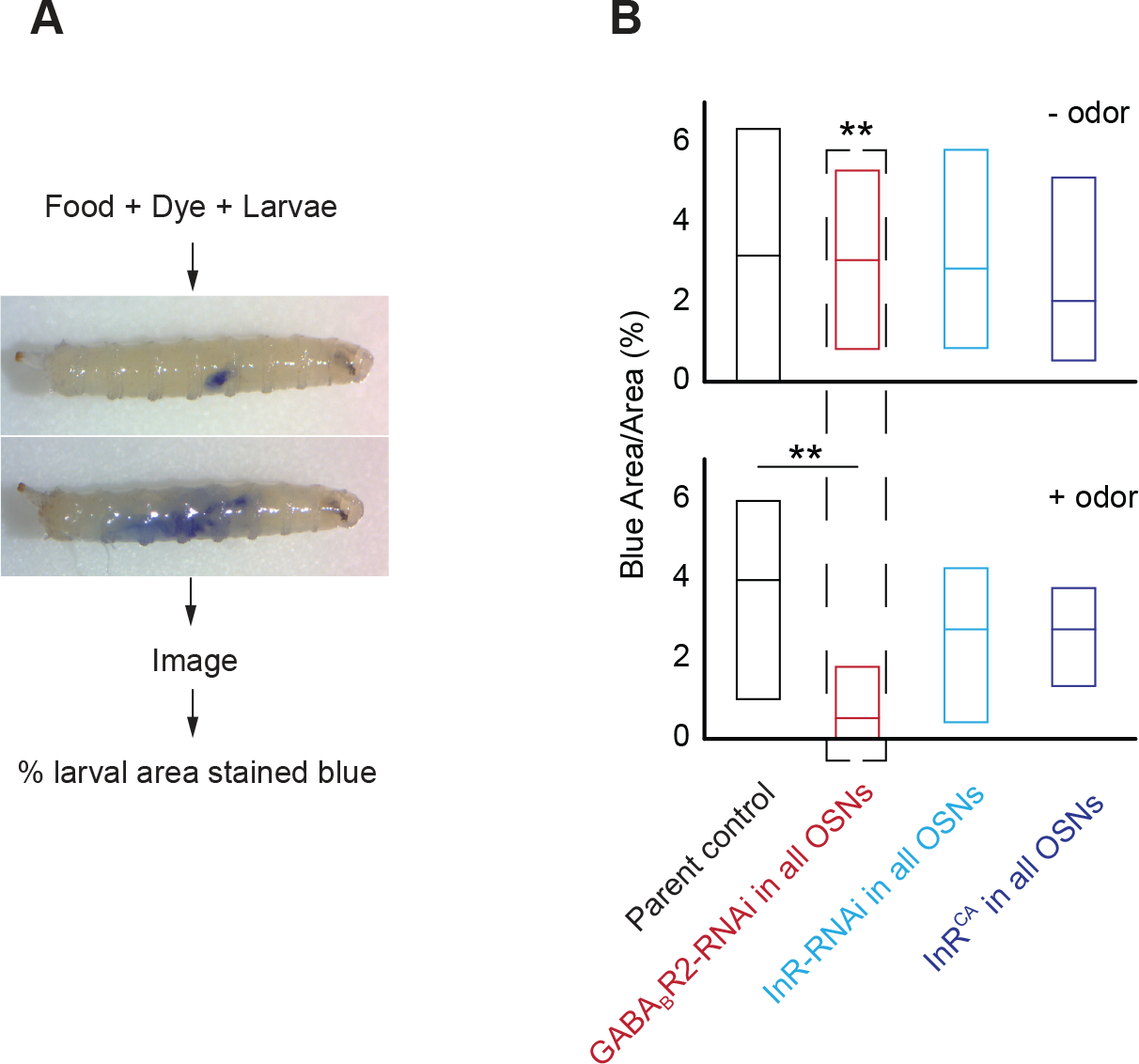
Larval food-intake measurements. (**A**) Schematic for food-intake assay. Larvae are allowed to feed on food mixed with a dye. After feeding, larvae are imaged, and food intake measured by counting the no. of pixels stained blue relative to the total no. of pixels in the whole larval body. (**B**) Data from the feeding assay in the absence (top panel) and presence (bottom panel) of odor (pentyl acetate, 10^−2^ vol:vol) are shown. Boxes are interquartile ranges. **p<0.001.

## RESULTS

### Starvation Enhances *Drosophila* Larval Olfactory Behavior Toward a Subset of Odors

Starvation increases olfactory sensitivity that encourage animals to search for food. This phenomenon has been demonstrated in several animals including humans (Cameron et al., 2012; Chao et al., 2004; Ko et al., 2015; Koelega, 1994; Root et al., 2011; Stafford & Welbeck, 2011; J. Tong et al., 2011). We asked whether starvation alters olfactory behavior in the *Drosophila* larva. We used the simple two-choice assay to measure larval behavior responses to an odor source (***Figure 1A***). In this assay, the larvae are offered a choice between an odor and a control diluent placed on filter paper discs on opposite sides of the arena. A response (attractive) index is measured based on number of larvae on either half of the plate (Monte et al., 1989; Rodrigues & Siddiqi, 1978). We measured behavior responses of starved or non-starved wild-type larvae to a panel of seven odorants using this assay. The odorants were selected based on their ability to elicit strong, specific physiological responses from one or few OSNs (2,5-dimethylpyrazine ∷ OSN33b; acetal ∷ OSN42b; pentyl acetate ∷ OSN47a; 4-hexen-3-one ∷ OSN42a; 4,5-dimethylthiazole ∷ OSN59a; anisole ∷ OSN30a; ethyl acetate ∷ OSN42a & OSN42b) (Kreher et al., 2008; Mathew et al., 2013). We noted that starved third-instar larvae were more attracted to odors than non-starved larvae. However, this increase in attraction was statistically significant in only three of the seven odors tested (Acetal: RI= 0.47 (Starved) vs. 0.33 (non-starved), *p<0.001*; pentyl acetate: RI= 0.56 (Starved) vs. 0.48 (non-starved), *p<0.05*; 4,5-dimethylthiazole: RI= 0.35 (Starved) vs. 0.21 (non-starved), *p<0.001*) (***Figure 1B***). These results suggest that starvation modulates larval olfactory function to increase attraction toward odors. The increase in attraction to only select odors suggests that starvation-dependent modulation may differentially impact individual neurons in the olfactory circuit.

### InR and GABA_B_R are Necessary for Starvation-Dependent Modulation of Olfactory Behavior

While starvation enhances larval behavior toward odors, the precise mechanisms by which the animal’s starved state modulates OSN function and olfactory behavior remain unclear. Insulin and GABA signaling have been implicated to play important roles during both starvation and olfactory behavior in insects and mammals (Avery & Horvitz, 1990; Enell et al., 2010; Li et al., 2015; Root et al., 2008; Sohrabipour et al., 2018; Wan et al., 1997; Weizman et al., 1990). Insulin and GABA signals are transduced via corresponding receptors (InR and GABA_B_-R respectively) expressed on neuronal membranes. While InR and GABA_B_-R expression have been clearly demonstrated at terminals of adult fruit fly OSNs (Ignell et al., 2009; Ko et al., 2015; Root et al., 2011), we wanted to confirm their expression at the terminals of larval OSNs. We used an anti-InR antibody to label InR in dissected third-instar larval preparations. We found that InR was expressed at the terminals as well as along the axonal projections of larval OSNs (***Figure 2A,B,C,D***). We generated an anti-GABA_B_-R1 antibody (*see materials and methods)* to characterize the distribution of GABA_B_-receptors in larval OSNs. We found that GABA_B_-R1 also localized to the terminals as well as axonal projections of larval OSNs (***Figure 2A,E,F,G***).

Next, we studied the roles of InR and GABA_B_-R in OSNs during starvation-dependent modulation of olfactory behavior. We used the UAS-Gal4 system to manipulate InR and GABA_B_-R levels in either all OSNs (using *Orco-Gal4*; (Larsson et al., 2004)) or in a single pair of OSNs (using *Or47a-Gal4*; (Vosshall et al., 2000)). *InR* levels were altered in OSNs by driving either a *UAS-InR-RNAi* (↓ InR levels) (Vienna Drosophila RNAi (VDRC) Stock Center) or a constitutively active version of *InR*: *UAS-InR^CA^* (↑ InR activity) (Root et al., 2011). GABA_B_R levels were reduced in OSNs by driving a *UAS-GABA_B_R2-RNAi* (Root et al., 2008). Since OSN∷47a responds strongly to pentyl acetate (Mathew et al., 2013), we measured larval attraction to pentyl acetate (10^−2^ vol:vol) in the two choice assay. The data are presented in ***Figure 2H***. Among the two control strains tested, starved larvae showed higher attraction to pentyl acetate than non-starved larvae. Reducing InR levels in either all OSNs or only in OSN∷47a did not alter this trend. However, increasing InR activity by driving the expression of *UAS-InR^CA^* impacted the starvation-dependent modulation of olfactory behavior. Starved larvae in which InR activity was increased in all OSNs showed a significantly reduced attraction to the pentyl acetate (RI = 0.28) compared to non-starved larvae (RI = 0.47; *p<0.05*). Even more dramatic effects were observed in both starved and non-starved larvae in which InR activity was increased in only OSN∷47a. Both non-starved larvae (RI = −0.15) and starved larvae (RI = −0.10) were repulsed by the odor. Reducing GABA_B_R levels also impacted the starvation-dependent modulation of olfactory behavior. While the overall attraction toward pentyl acetate was lower in all cases, reducing GABA_B_R levels in all OSNs eliminated the starvation-dependent increase in attraction observed in control animals (*p>0.05*). A more dramatic effect was observed in starved larvae when GABA_B_R levels were reduced in only OSN∷47a. In this case, the non-starved larvae were attracted to the odor (RI = 0.15) while the starved larvae were repulsed by the odor (RI = −0.14; *p<0.05*).

These results suggest that both insulin signaling and GABA signaling are necessary for starvation-dependent modulation of OSN function. Consistent with the prevailing model of OSN modulation, our results suggest that reducing insulin signaling in OSNs mimics starved conditions and thus the starvation-dependent increase in attraction to odor remains intact. On the other hand, increasing insulin signaling artificially mimics non-starved conditions and thereby reduces attraction to odor. The role of GABA signaling in this mechanism was previously unexplored. Our results suggest that it plays an opposite role compared to insulin signaling, in that lower GABA signaling possibly mimics non-starved conditions for the animal.

### GABA_B_R and InR Expression Levels are Sensitive to the Animal’s Starved State

Since manipulating InR and GABA_B_R levels in OSNs impacted starvation-dependent modulation of OSN function, we wondered whether their expression levels in larval CNS are sensitive to the animal’s starved state. To test this, we dissected heads of starved and non-starved larval samples, extracted mRNA from the dissected heads and carried out gene expression analyses using RT-qPCR (***Figure 3A***). We evaluated the relative gene expression of genes involved in the insulin and GABA signaling pathways. The data are presented in ***Figure 3B***&***C***. For genes involved in the insulin signaling pathway, we tested expression levels of InR as well as eight *Drosophila* Insulin-Like Peptides (DILPs) (four shown here) (Geminard et al., 2006). Consistent with the prevailing model of OSN modulation, we noted that starvation decreased expression levels of several insulin signaling components including InR (20% decrease, *p<0.05*), DILP-2 (22% decrease, *p<0.05*), and DILP-6 (40% decrease, *p<0.001*) (***Figure 3B***). For genes involved in the GABA signaling pathway, we tested the expression levels of Glutamate decarboxylase (GAD1), an enzyme responsible for catalyzing the production of GABA, GABA_A_R and GABA_B_R subunits (Bettler et al., 2004). We noted that starvation decreased expression levels of one of the three *GABA_B_-receptor* subunits, *GABA*_*B*_*R2* (25% decrease, *p<0.05*) but increased the expression of *GABA*_*A*_*R* (45% increase, *p<0.05*). These results suggest that expression levels of insulin and GABA signaling components in the larval CNS are sensitive to the animal’s starved state.

### Novel Method to Evaluate Gene Expression Levels in Larval OSNs

Although the above gene expression analysis in larval heads provided information consistent with the prevailing OSN modulation model, we wanted to evaluate gene expression levels specifically in larval OSNs, especially in the context of high or low insulin signaling. However, evaluating gene expression changes specifically in OSNs posed a technical challenge. To overcome this challenge and to carry out OSN-specific gene expression studies, we adapted a previously established protocol to isolate single-cell type nuclei (Ma & Weake, 2014). Using this technique, we successfully isolated larval OSN nuclei and carried out OSN-specific gene expression analyses. Briefly, OSN nuclei were genetically tagged using a *UAS-eGFP-Msp300*^*KASH*^ construct (containing a localization signal for the nuclear membrane) (***Figure 4A, B***). GFP-tagged nuclei were separated and enriched using an affinity-based pull-down approach (***Figure 4C***). RNA extracted from enriched OSN nuclei were then used as substrate for gene expression analysis. We validated the effectiveness of OSN isolation by comparing relative gene expression levels from pre-isolation nuclei and post-isolation nuclei samples. Significant enrichment of OSN specific genes such as *Orco* (>2.5 fold; *p*<*10*^−*6*^) and *Or94b* (>4.5 fold; *p*<*10*^−*6*^) and eGFP (>2.0 fold; *p*<*1.5* × *10*^−*4*^) but not genes such as *Syt1* and *Act42a* that are commonly expressed in most neuron types support the effectiveness of OSN isolation using this technique (***Figure 4D***). Successful implementation of this technique in our lab enables us to isolate and enrich OSN nuclei from the thousands of heterogeneous cell-type nuclei in the *Drosophila* larva, extract RNA from the isolated nuclei, and compare gene expression analyses specifically in OSNs. We used this technique below to evaluate OSN gene expression under conditions of high and low insulin signaling.

### Insulin and GABA Signaling Pathways Interact in Larval OSNs

While several studies have suggested that insulin and GABA signaling pathways interact to mediate neuronal function in insects and mammals (Enell et al., 2010; Sohrabipour et al., 2018; Wan et al., 1997), it was not clear whether these signaling pathways interact within OSNs to mediate OSN function. To test any potential interaction between the signaling pathways, we evaluated gene expression in OSNs using the method implemented in our lab (***Figure 4***). We predicted that if the two signaling pathways interact in OSNs, then affecting one pathway would impact the other. We generated two separate experimental lines, each expressing *UAS-eGFP-Msp300*^*KASH*^ along with either *UAS-InR-RNAi* (↓ InR levels) or *UAS-InR^CA^* (↑ InR activity) in all OSNs. We enriched GFP-tagged OSN nuclei from control larvae and the two experimental larval strains. We extracted RNA from the isolated OSN nuclei and quantified expression levels of InR and the three GABA_B_R subunit genes (***Figure 5A***). Normalized gene expression levels are plotted in ***Figure 5B***&***C***. We found that decreasing InR levels significantly reduced expression levels of *GABA*_*B*_*R1* (54% decrease, *p<0.001*) and *GABA*_*B*_*R2* (25% decrease, *p<0.05*) subunits while increasing InR levels significantly increased expression levels of *GABA*_*B*_*R2* (35% increase, *p<0.05*) and *GABA*_*B*_*R3* (90% increase, *p<0.05*) subunits (***Figure 5B***). These results suggest that GABA and insulin signaling pathways interact within OSNs, with GABA_B_R activity potentially downstream of InR activity.

### Potential Downstream Targets of Insulin Signaling in Larval OSNs

The above result, which suggests that InR levels in OSNs impact expression of GABA_B_R subunit genes is consistent with recent studies claiming that cell-surface InR translocates to nucleus, associates with promoters, and regulates gene expression (Hancock et al., 2019). Therefore, we wondered whether altering InR levels specifically in OSNs altered expression of other downstream genes known to play a role in olfaction. To test this, we compared relative expression levels of known OSN genes in larval strains that were manipulated to have high or low InR activity in OSNs.

We noted that decreasing insulin signaling in OSNs led to ∼50% increase in *sNPFR1* expression levels in OSNs (*p<0.05*) (***Figure 5B***). This result is consistent with predictions of the prevailing model, which claims that low insulin signaling during the animal’s starved state leads to increased levels of *sNPFR1*, which in turn enhances OSN facilitation and response to odors (Bargmann, 2012; Ko et al., 2015). Next, we looked at the impact of InR activity on six other OSN specific genes (***Figure 5B***). We noted that decreasing insulin signaling in OSNs significantly decreased expression of olfaction genes such as *Orco* (61% decrease; *p<0.001*), *Rutabaga (Rut)* (24% decrease; *p*<*0.05*), and *Synaptotagmin (Syt1)* (24% decrease; *p*<*0.05*) compared to wild type controls. *Orco* is the co-receptor for odor receptors expressed in OSNs (Sato et al., 2008; Smart et al., 2008; Wicher et al., 2008). *Rut* encodes adenylyl cyclase, a component of olfactory signal transduction (Cho et al., 2004). *Syt1* plays a role in neurotransmission (Kidokoro, 2003). However, increasing or decreasing insulin signaling had no impact on other OSN genes tested including two other components of olfactory signal transduction, *G_αi_* and *G_olf_* (Boto et al., 2010; Kaupp, 2010). While this is not an exhaustive list of OSN genes, our results so far suggest that downstream targets of insulin signaling in OSNs potentially play important roles in odor detection, olfactory information processing, and neurotransmission.

### GABA_B_R and InR Levels in OSNs Impact Larval Body Weight

Since an inability to regulate sensitivity to food odors at appropriate times could lead to irregular foraging habits, which in turn could impact weight gain, we wondered whether altering insulin and GABA signaling specifically in the OSNs would affect the animal’s overall body weight. In support of this warrant, recent studies have shown that genetically obese rats have low levels of insulin in the brain including the olfactory bulb and imbalanced insulin signaling via insulin receptors is associated with obesity phenotypes (Baskin et al., 1985; Kubota et al., 2017).

We conducted a careful analysis of *Drosophila* larval body weight in genotypes expressing high or low levels of InR or GABA_B_R in the OSNs. We found that altering *GABA*_*B*_*R* or *InR* levels in OSNs led to significant increases in larval body-weight compared to parental control (***Figure 6***). This result reveals an interesting link between OSN regulatory mechanisms and animal physiology.

### GABA_B_R Levels in OSNs Impact Larval Feeding Behavior

Since altering *GABA*_*B*_*R* or *InR* levels in OSNs led to increases in body weight, we hypothesized that the body weight increases are due to altered food consumption in mutant genotypes. We used a larval feeding (food + dye intake) assay to test food consumption in larval genotypes in which *GABA*_*B*_*R* or *InR* levels were altered in OSNs (***Figure 7A***) (Kaun et al., 2007). We found that in the absence of any odor, wild type larvae and larvae expressing altered levels of *GABA*_*B*_*R* and *InR* in OSNs have similar levels of food intake in a 15 min period (***Figure 7B***, *Top panel*). A previous study demonstrated that larvae engage in appetitive cue-driven feeding behavior (Y. H. Wang et al., 2013). When the assay was conducted in the presence of a food odor like pentyl acetate, we found that larvae expressing GABA_B_R-RNAi had 70% lower levels of food intake compared to other genotypes (*p*<*0.001*) (***Figure 7B***, *Bottom panel*). These results suggest that manipulating OSN modulation mechanisms not only impact foraging behaviors (***Figure 2B***) but also feeding behaviors. While a decrease in food consumption in a 15 min period may not explain the increase in body weight in genotypes in which *GABA*_*B*_*R* levels were decreased in OSNs, it is possible that other aspects of feeding behavior such as frequency of feeding, which is difficult to test in larvae, may also be affected.

## DISCUSSION

Starvation dependent increase in larval behavior toward odors (***Figure 1***) requires both insulin and GABA signaling in OSNs (***Figure 2B***). Insulin and GABA signaling pathways interact within OSNs (***Figure 5A***) and likely modulate OSN function by impacting odor reception (*Orco*), olfactory information processing (*Rut*), and/or neurotransmission (*Syt1*) (***Figure 5B***). Defects in GABA/insulin signaling pathways impact the animal’s feeding behavior and body weight (***Figures 6 and 7***). These findings suggest a hitherto unsuspected role for GABA signaling in starvation-dependent modulation of OSN function, a role that is likely downstream of insulin signaling. They also raise questions about how individual OSNs may be differentially modulated by the animal’s starved state. Finally, these findings imply a potential relationship between nutrient sensing and animal physiology. Overall, this study challenges the prevailing mechanistic model of starvation dependent modulation of OSNs. Results from this study will enable the generation of new hypotheses that will have significant implications for understanding general principles and mechanisms by which an animal’s internal state modulates sensory function.

### Interactions between Insulin and GABA Signaling Pathways

GABA and insulin signaling play important roles during both starvation and olfactory behavior (Avery & Horvitz, 1990; Enell et al., 2010; Li et al., 2015; Root et al., 2008; Sohrabipour et al., 2018; Wan et al., 1997; Weizman et al., 1990). While GABA signaling in different regions of the animal brain is known to mediate starvation-dependent behavior, its role in specific olfactory neurons during starvation is unclear (Avery & Horvitz, 1990; Weizman et al., 1990). Similarly, insulin has long been considered as an important mediator of state dependent modulation of feeding behavior. However, its precise role in olfactory neurons during starvation is controversial. According to the prevailing model, insulin signaling decreases upon starvation (Bargmann, 2012; Ko et al., 2015). However, a previous study showed that there is a three-fold increase in *DILP-6* (*Drosophila* Insulin like Peptide) mRNA expression in larval tissue including fat bodies upon starvation, which is inconsistent with the model (Slaidina et al., 2009). While the significance of *DILP-6* increase in larval tissue during starvation is as yet unclear, consistent with the prevailing model, we show that InR and *DILP-6* expression in larval head samples decrease upon starvation (***Figure 3B***).

We also show that higher insulin signaling increases expression levels of GABA_B_Rs in OSNs (***Figure 5B***). This result is in line with several other studies in flies and mammals that have suggested possible interactions between GABA signaling and insulin signaling in different regions of the brain. The most relevant example supporting our observation is noted in mice where insulin increases the expression of GABA_A_Rs on the postsynaptic and dendritic membranes of CNS neurons (Wan et al., 1997). Other examples show how GABA signaling might influence insulin signaling. For instance, in flies, GABA signaling from interneurons has been shown to affect insulin signaling by regulating DILP production (Enell et al. 2010); In humans, GABA administration significantly increases circulating insulin levels under both fasting and fed conditions (Li et al., 2015; Sohrabipour et al., 2018); In diabetic rodent models, combined oral administration of GABA and an anti-diabetic drug (Sitagliptin) promoted beta cell regeneration and reduced blood glucose levels (Liu et al. 2017; Sohrabipour et al. 2018). Overall, our study adds to this growing body of literature and strongly suggests that GABA and insulin signaling pathways interact within larval OSNs to mediate OSN modulation.

### Differential Modulation of OSN Function

We note that starvation enhanced larval attraction toward only a subset of the odors tested (***Figure 1***). An intriguing question in the field is whether starvation enhances an animal’s ability to detect food-odors or all odors. Studies are inconclusive so far. Some studies have shown that starvation enhances an animal’s ability to detect both food-related odors (Apelbaum et al., 2005) and nonfood-related odors (Aime et al., 2007). While similar results have also been shown in humans, the findings regarding the relevance of odor to feeding are rather mixed (Koelega, 1994; Stafford & Welbeck, 2011). This study along with previous studies from our lab and others raise the possibility that starvation differentially modulates individual OSNs. Indeed, individual OSNs exhibit functional diversity that may lend them to differential modulation by the animal’s internal state (Clark et al., 2018; Newquist et al., 2016; Slankster et al., 2019). This diversity may stem from heterogeneous GABA_B_R levels on the terminals of individual OSNs that determine differential presynaptic gain control (Root et al., 2008). It is reasonable to speculate that heterogeneous GABA_B_R and/or InR levels in individual OSNs could contribute to differential modulation of OSNs by the animal’s starved state, which in turn impacts behavior toward only a subset of odors.

### Sensory Neuron Modulation and its Impact on Feeding Behavior and Weight Gain

An inability to regulate sensitivity to food odors at appropriate times leads to irregular feeding habits, which in turn leads to weight gain. Obesity researchers will readily acknowledge that while several obvious risk factors for obesity (e.g. genetics, nutrition, metabolism, environment etc.) have been heavily researched, the relationship between nutrient sensing/sensory behavior and obesity remains grossly understudied. The present study sets the stage to further explore this relationship. Interestingly, several of the signaling molecules described in this study that play a role in OSN modulation have also been implicated in hyperphagia and obesity phenotypes. For instance, overexpression of *sNPF* in *Drosophila* and NPY injection in the hypothalamus of rats leads to increased food-intake and bigger and heavier phenotypes (Lee et al., 2004; Wahlestedt et al., 1993). Genetically obese rats have low levels of insulin in the brain including in the olfactory bulb and imbalanced insulin signaling via insulin receptors is associated with obesity phenotypes (Baskin et al., 1985; Kubota et al., 2017). Adenylyl cyclase (*rut*) deficient mice were found to be obese (Z. S. Wang et al., 2009) and both *Adenylyl cyclase3* and *Synaptotagmin4* have been targeted for anti-obesity drug development (Q. C. Tong, 2011; Wu et al., 2016). These studies provide added significance to our observation that manipulating mechanisms mediating starvation-dependent modulation of OSNs impact feeding behavior and weight gain in larvae.

Indeed, food odors can be powerful appetitive cues. A previous study showed that larvae engage in appetitive cue-driven feeding behavior and that this behavior required NPF signaling within dopaminergic neurons in higher-order olfactory processing centers (Y. H. Wang et al., 2013). Our studies show that manipulating GABA_B_R signaling in first-order OSNs impact appetitive cue-driven feeding behavior in larvae. While it remains to be seen whether parallel regulations during different stages of olfactory information processing impact feeding behavior, further studies are needed to reveal the mechanistic relationship between GABA_B_R/InR signaling in OSNs, feeding behavior, and changes in body-weight.

### Motivating Model for Future Investigations

Based on the evidence so far, we propose a motivating model for future investigations (***Figure 8***). In this model, GABA_B_R and InR expressed on the terminals of larval OSNs act as sensors for the internal state of the animal. Their concerted activity impacts OSN function either at the level of odor reception by affecting the expression of *Orco* or at the level of olfactory signal transduction by affecting the expression of *Rut* or at the level of neurotransmission by affecting the expression of *Syt1* and *sNPF*. We acknowledge that more exhaustive gene expression analyses are required to identify other molecular players downstream of InR and GABA_B_R. It would also be valuable to investigate the relationship between InR expression levels on the terminals of individual OSNs and the sensitivity of individual OSNs to modulation by the animal’s starved state.

**Figure 8.**
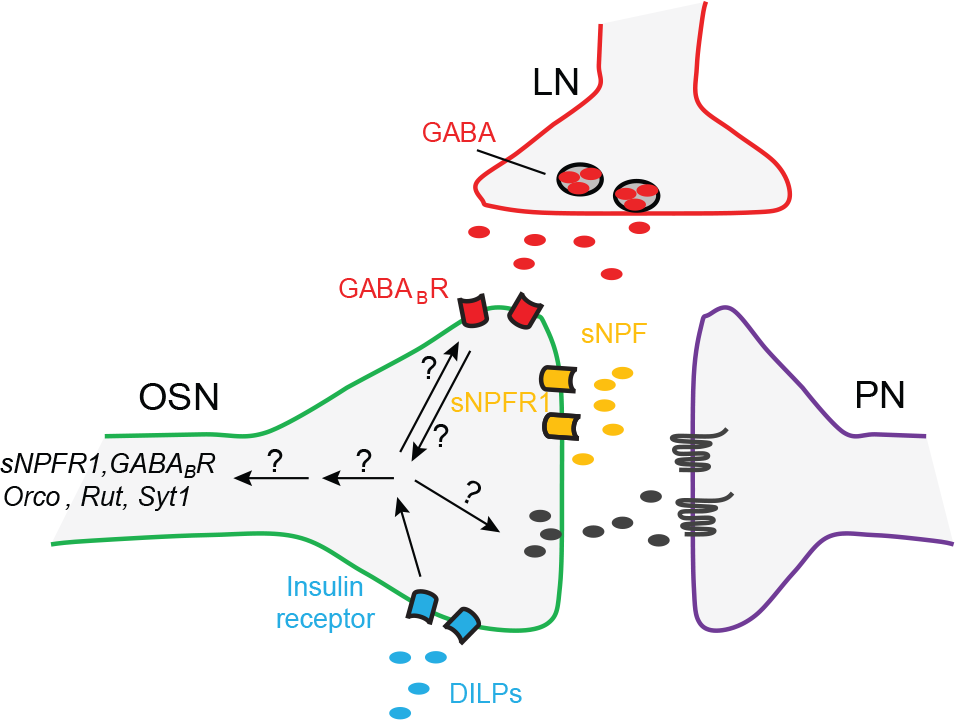
Updated model for OSN modulation. A cartoon model of neuronal interactions in the olfactory glomerulus of the larval antennal lobe and OSN-modulation mechanisms is shown. In a glomerulus, OSNs make synaptic connections with second-order projection neurons (PNs) and local neurons (LNs). Within OSNs, insulin signaling interacts with GABA signaling to impact OSN function. Insulin signaling potentially impacts the expression of downstream genes including *sNPFR1*, *GABA*_*B*_*R*, *Orco*, *Rut*, and *Syt1*. Based on published data from other labs as well as data from this study, during the animal’s starved state, low insulin signaling leads to higher sNPFR1 levels, which in turn lead to increases in OSN facilitation and response to attractive odors. However, the precise mechanisms downstream of GABA_B_R/InR receptor activity that effect changes in gene expression and OSN facilitation remain unclear.

### Potential Caveats of this Study

A valid concern is that an innate attraction of larvae toward an odorant does not necessarily equate to food-search behavior. However, we argue that attractiveness toward an odor source is a reliable measure of food-search behavior because an animal’s ability to efficiently smell and move toward an odor source necessarily predicates most forms of such behavior (Gershow et al., 2012; Gomez-Marin et al., 2011; Pierce-Shimomura et al., 1999; Sourjik & Wingreen, 2012). Another possibility to be considered is that changes in OSN sensitivity, food-search and/or feeding behaviors are independently regulated. For instance, Yu and colleagues noted that starvation-induced hyperactivity in adult *Drosophila* was independently regulated from food consumption behavior in the flies. They showed that blocking octopamine signaling in a small group of octopaminergic neurons located in the subesophageal zone (SEZ) of the fly brain neurons eliminated starvation induced hyperactivity but not the increase in food consumption (Yu et al., 2016). While we cannot rule out such a possibility, the evidence presented in this study support the argument that starvation induced-changes in OSN function is related to the observed changes in food search and feeding behaviors. Finally, while we tested the hypothesis that increases in body-weight of mutant genotypes are due to altered food consumption, we have not yet tested alternate hypotheses that body-weight increases may be due to altered metabolism or increased fat accumulation.

### General Impacts of this Study

Our study conducted in a simple, tractable, and highly conserved model system challenges the prevailing model of starved-state dependent modulation of OSN function. It highlights and offers unique opportunities that are now possible to address our inadequate understanding of OSN modulation mechanisms at the resolution of single neurons, which in turn would help us better understand how flexibility and the ability to adapt to a particular internal state are built into the sensory circuit.

## Acknowledgments

We are grateful to Samantha Badie for technical assistance. Research reported in this publication was supported by Startup funds from the University of Nevada, Reno and by a grant from the NIGMS of the National Institute of Health under grant number P20 GM103650 to DM. Research reported here used the Cellular and Molecular Imaging Core facility supported by the National Institute of General Medical Sciences of the National Institutes of Health under grant number P20 GM103650.

---

**Supplementary table 1.**
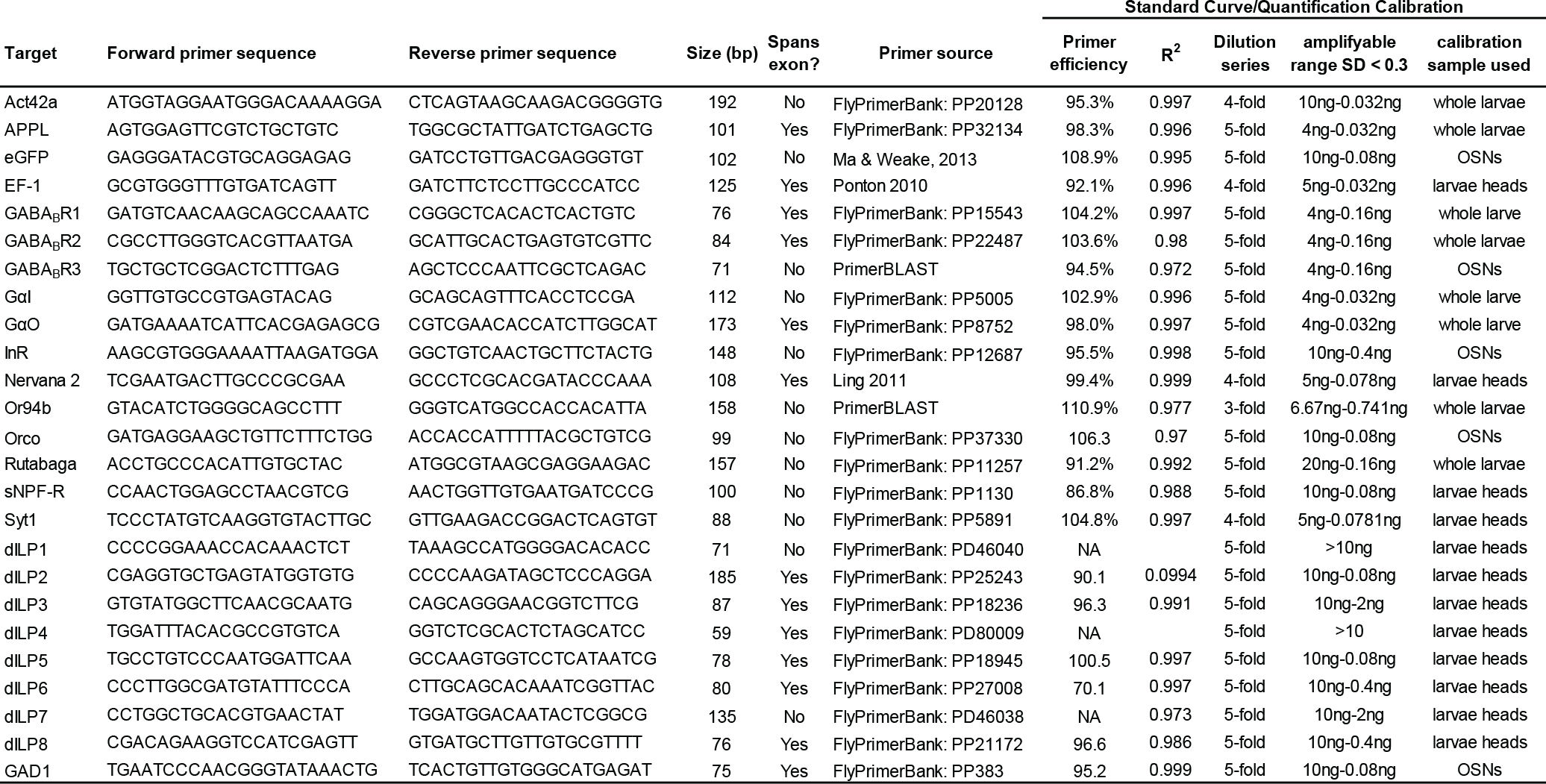
Primers used in RT-qPCR analysis. Forward and reverse primer sequences used in RT-qPCR analyses, their source, and calculated primer efficiencies are shown. Expected sizes of RT-PCR products were estimated using PrimerBLAST (https://www.ncbi.nlm.nih.gov/tools/primer-blast/).

## Notes

**Conflict of Interest:** The authors declare no competing financial interests.

